# Ion Identity Molecular Networking in the GNPS Environment

**DOI:** 10.1101/2020.05.11.088948

**Authors:** Robin Schmid, Daniel Petras, Louis-Félix Nothias, Mingxun Wang, Allegra T. Aron, Annika Jagels, Hiroshi Tsugawa, Johannes Rainer, Mar Garcia-Aloy, Kai Dührkop, Ansgar Korf, Tomáš Pluskal, Zdeněk Kameník, Alan K. Jarmusch, Andrés Mauricio Caraballo-Rodríguez, Kelly Weldon, Melissa Nothias-Esposito, Alexander A. Aksenov, Anelize Bauermeister, Andrea Albarracin Orio, Carlismari O. Grundmann, Fernando Vargas, Irina Koester, Julia M. Gauglitz, Emily C. Gentry, Yannick Hövelmann, Svetlana A. Kalinina, Matthew A. Pendergraft, Morgan W. Panitchpakdi, Richard Tehan, Audrey Le Gouellec, Gajender Aleti, Helena Mannochio Russo, Birgit Arndt, Florian Hübner, Heiko Hayen, Hui Zhi, Manuela Raffatellu, Kimberly A. Prather, Lihini I. Aluwihare, Sebastian Böcker, Kerry L. McPhail, Hans-Ulrich Humpf, Uwe Karst, Pieter C. Dorrestein

**Author notes:** These authors contributed equally (Robin Schmid, Daniel Petras, Louis-Félix Nothias). **Author contributions.** **General conceptualization** RS, DP, LFN, MW, PCD conceptualized the idea of IIMN and its integration into GNPS and feature-finding software tools RS, DP, LFN, PCD wrote the manuscript RS, BA, FH, HUH conceptualized the MZmine feature grouping workflow UK, HH provided discussion and feedback on IIMN and the MZmine workflow **Development** RS developed the IIMN modules in MZmine and the MS^2^ spectral library generation modules MW, RS developed the “supplementary edges” format in the FBMN workflow to enable IIMN MW programmed the IIMN workflow on GNPS RS, MW developed the direct submission of MZmine data to run IIMN on GNPS JR, MGA developed the XCMS/CAMERA IIMN integration in R HT developed the MS-DIAL FBMN and IIMN integration KD developed the MS^2^ spectral merge function into the export modules for FBMN, IIMN, and SIRIUS, which was coordinated by SB TP, AK provided feedback and help for the development and integration of IIMN in MZmine **Experiments, data analysis, validation** DP, LFN, AA, AAO, GA, AB, ATA, AMCR, JMG, ECG, COG, YH, ANJ, AKJ, SK, ZK, IK, ALG, KLM, MNE, MAP, MWP, RT, FV, KW performed experiments, analyzed data with the MZmine IIMN workflow, made data publicly available through MassIVE, and validated the results. KAP, MR, HZ, HUH, PCD provided data and resources RS, DP, ATA, ANJ analyzed data ATA, RS, ANJ wrote supplemental use cases YH, SK, ANJ, AK, BA, ZK tested and provided feedback on the MZmine workflow **Documentation and videos** LFN, HMR, AB, DP, MW, ATA, RS, MNE created the IIMN and FBMN documentations RS produced video tutorials on FBMN, IIMN, and MZmine MW, RS produced videos on FBMN and the direct submission of MZmine results to GNPS DP, MW produced a video tutorial for feature finding with MZmine and FBMN in GNPS All authors contributed to the final manuscript.

## Abstract

Molecular networking connects tandem mass spectra of molecules based on the similarity of their fragmentation patterns. However, during ionization, molecules commonly form multiple ion species with different fragmentation behavior. To connect ion species of the same molecule, we developed Ion Identity Molecular Networking. These new relationships improve network connectivity, are shown to reveal novel ion-ligand complexes, enhance annotation within molecular networks, and facilitate the expansion of spectral libraries.

## Main

Molecular networking (MN)^1^ within the GNPS web platform (http://gnps.ucsd.edu)^2^ has been used for the analysis of non-targeted mass spectrometry data in various fields^3,4^ MN relies on the principle that similar structures tend to form similar patterns in tandem mass spectra (MS^2^). By computing pairwise spectral comparisons in a dataset, we create an MS^2^ spectral network. This network is enriched by annotating the MS^2^ spectra against MS^2^ spectral libraries or compound databases (Figure 1); further, annotations can be propagated through the network^5^. MN can be used to map the chemical space of complex samples to facilitate the discovery of new molecules, especially analogs of known compounds^2^. For the data analysis of liquid chromatography-mass spectrometry (LC-MS^2^) data, feature-based molecular networking (FBMN) combines MN with chromatographic feature finding tools^6^. During LC-MS ionization, a given compound can generate multiple ion adducts *(e.g.,* protonated and sodiated), which appear as individual nodes in a molecular network, due to different precursor mass-to-charge ratios *(m/z).* As various commonly detected ion adducts exhibit different fragmentation behavior during collision-induced dissociation (CID) (Supplementary Figure 1), MS^2^ spectral networking on its own might not connect any ion adducts of the same compound. Here, we present ion identity molecular networking (IIMN), a workflow that annotates and connects related ion species in feature-based molecular networks within the GNPS web platform.

**Figure 1:**
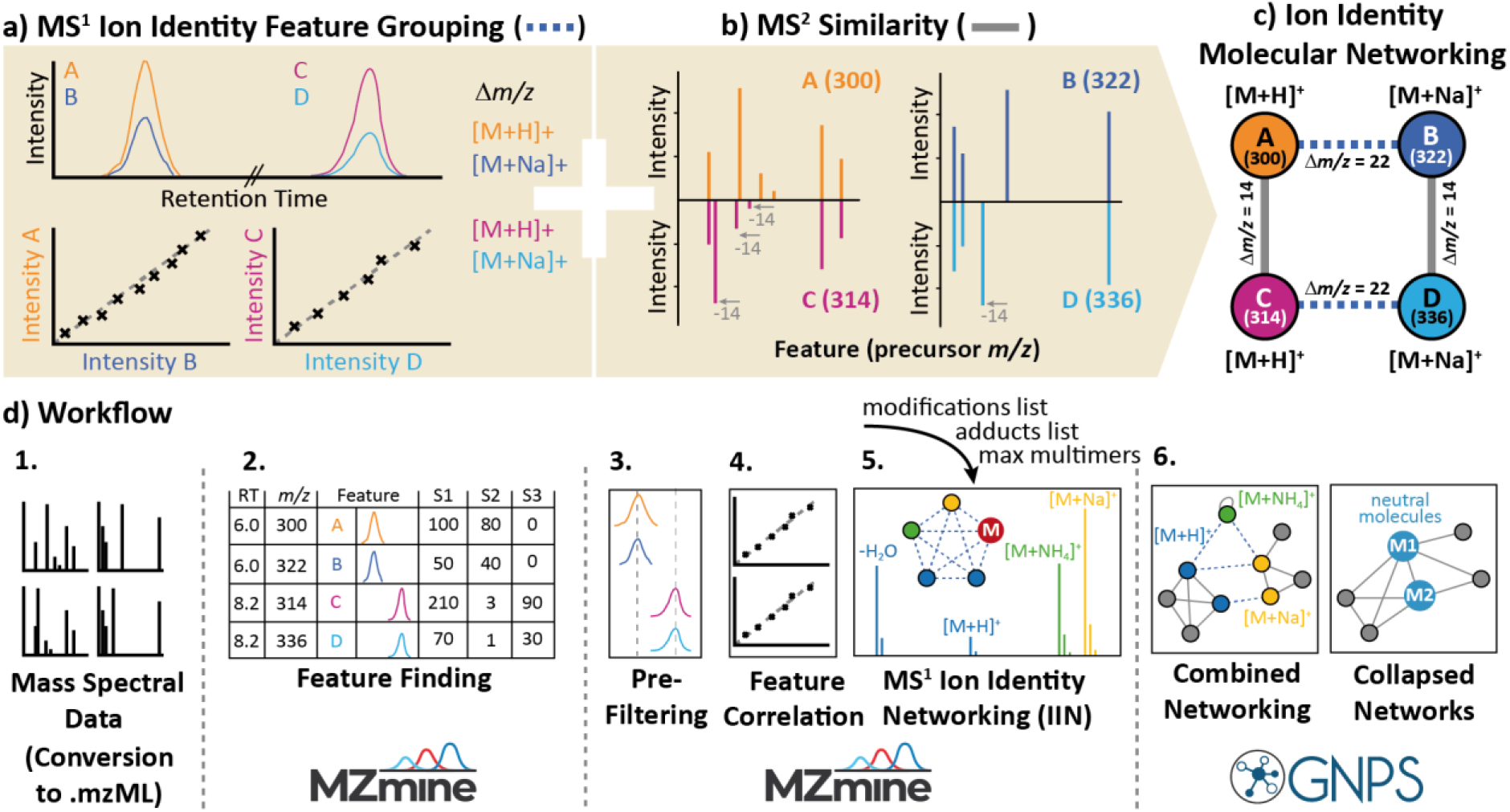
The concept of ion identity molecular networking (IIMN). **a**) shows the two main principles of the combined networks. IIN identifies and connects different ion species of the same compound based on MS^1^ characteristics, while FBMN connects LC-MS feature nodes by their MS^2^ fragmentation spectral similarity. **b)** highlights the data processing workflow to create combined IIMN networks in MZmine and GNPS. After feature detection and alignment across multiple samples, features are grouped based on the correlation of their chromatographic peak shapes and other MS^1^ characteristics. Subsequently, ion species of grouped features are identified with an ion identity library, which is generated based on user input for included adducts, in-source modifications, and a maximum multimers parameter. After uploading these results to GNPS, combined ion identity molecular networks are created on the webserver. Optionally, ion identity networks can be collapsed into single molecular nodes to reduce complexity and redundancy.

Multiple tools have been developed for the connection of ion species in LC-MS data, which typically compare retention time, chromatographic shape, and feature intensity across samples to group LC-MS features of the same compound^7–11^. Subsequently, ion species can be identified based on known mass differences^7^, resulting in MS^1^-based ion identity networks (IIN). We fully integrated IIN into MS^2^-based molecular networks in the GNPS environment. After feature grouping and identification of ion species, extracted data are uploaded to GNPS to run IIMN on the webserver. Resulting ion identity molecular networks contain two layers of feature (node) connectivity, linking ion identities of the same compound by MS^1^ characteristics and structurally similar compounds by MS^2^ spectral similarity (Figure 1). The IIMN modules in MZmine (Supplementary Figure 2) include new feature grouping and ion identity networking algorithms, as well as modules to visualize and analyze networking results.

To validate the identification of ion species with IIMN, we created an LC-MS^2^ benchmark dataset of a natural product mixture containing 300 compounds, in which we promoted adduct formation by post-column infusion of ammonium acetate or sodium acetate at different concentrations (Figure 2a-e). The IIMN networks can be depicted in alternative layouts that illustrate complementary results within the same dataset. It is also possible to collapse ion identity networks to reduce the redundancy of different ion species by merging them into a single “neutral molecule” (M) node (Figure 2c). In this dataset, IIN successfully connects ion identities and reduces the size of a complex network by 56% to four major compounds. The increased connectivity facilitates the propagation of structure annotations to neighboring in-source fragments and an unannotated compound. Finally, the abundance change of identified adducts ([M+H]^+^, [M+NH_4_]^+^, [M+Na]^+^) in our benchmark dataset is in agreement with the different postcolumn salt infusion conditions (H_2_O, NaAcetate or NH_4_ Acetate, Figure 2f) which validates ion species identification on a dataset level.

**Figure 2:**
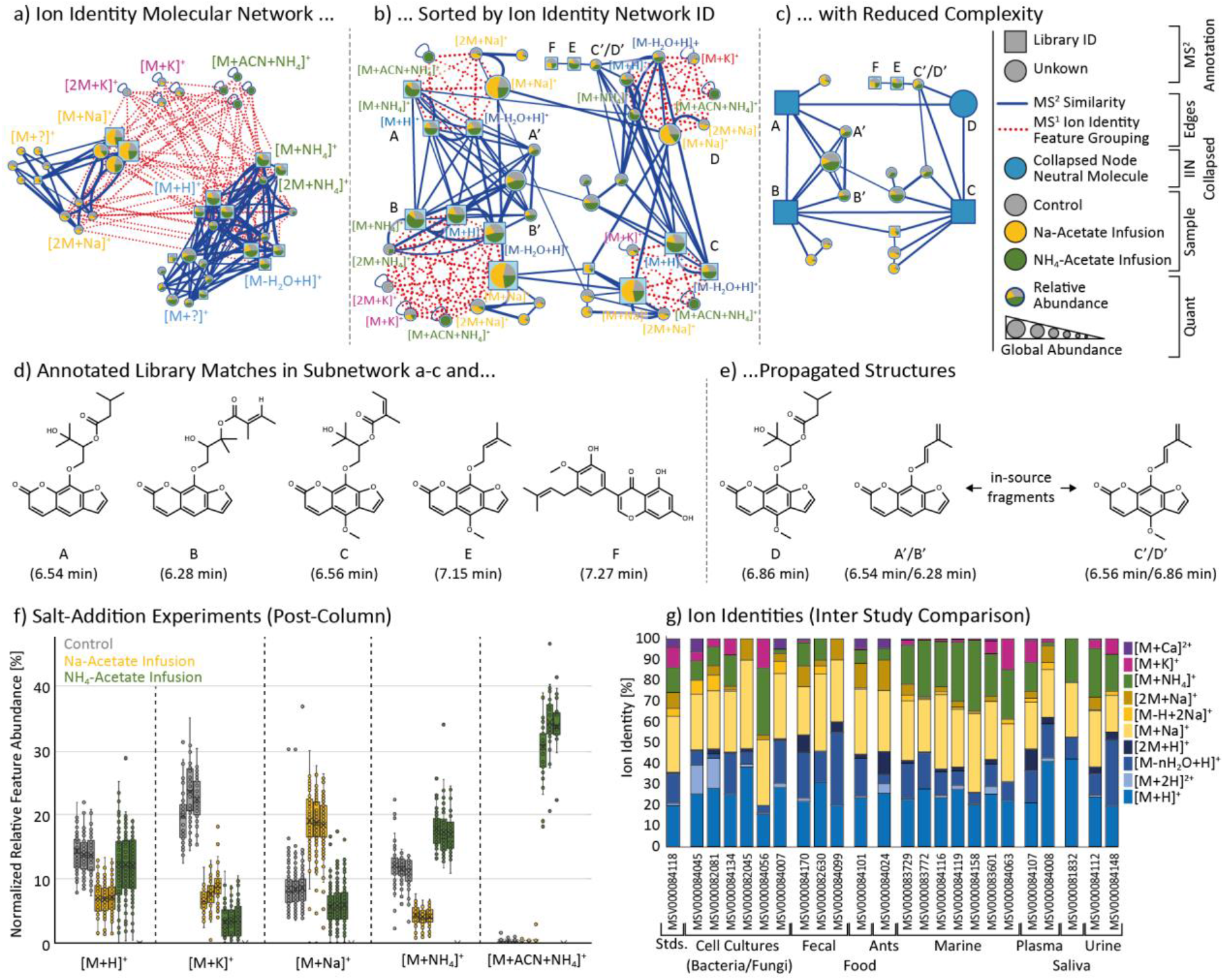
Ion identity molecular networking and statistical results. Depicted are three visualizations of the same ion identity molecular network from the post-column salt infusion experiments. **a**) Sorting by ion identities reveals that MS^2^ similarity edges often link sodiated ions ([M+Na]^+^ and [2M+Na]^+^) into a subnetwork that is separated from a subnetwork of ammonium adducts with protonated species. The pie-charts indicate relative abundances in different salt addition experiments (Control (H_2_O), grey; Na-Acetate, yellow, NH4-Acetate, green). The complexity and redundancy are reduced by **b**) sorting all ions of the same molecule in a circular layout and **c)** collapsing all IINs into single molecular nodes. This option reduces the complexity of this IIMN from 43 feature nodes to four molecular nodes (A-D) and 15 feature nodes (−56%). **d**) lists the structure of all GNPS library matches and **e)** propagated structures for D (based on A and C) and the in-source fragments A’ to D’. This subset of structurally related compounds gives a first statistical proof for high correct annotation rates during IIN in MZmine as adduct formation responds to the corresponding salt infusion, e.g., higher [M+Na]^+^ abundances in the sodium acetate buffer infusion. Moreover, this is also true on **f)** a dataset scale where the relative intensities of selected ion identities are plotted for each post-column infusion in triplicate. This plot reveals that the in-source cluster [M+ACN+NH_4_]^+^ exclusively forms in the ammonium acetate buffer infusion. **g**) IIMN results for 24 experimental datasets, showing the relative ion formation tendencies measured as the number of ion identities.

To test the workflow with data generated from various sample types and on different experimental platforms, 24 public datasets were processed by different authors using the MZmine workflow (Figure 2g, Supplementary Table 1). Here, the application of IIMN to identify post-column induced ion species can be particularly useful for the screening of biologically-relevant metal-binding compounds. In a native ESI-based metabolomics study, IIMN specifically revealed that the known siderophore yersiniabactin also acts as a zincophore (Supplementary Figure 3)^12^. In a dataset with 88 animal bile acid extracts, multiple smaller networks and unconnected nodes were combined to a large network of free bile acids and those conjugated to amino acids or sulfate, resulting in higher connectivity in the network (Supplementary Figure 4). IIMN also yielded additional structural information in the case of mold samples from *Stachybotrys chartarum* (Supplementary Figure 5). The increasing number of aliphatic hydroxyl groups in phenylspirodrimane derivatives was reflected by the maximum number of in-source water losses, whereas acetylation of hydroxy groups reduced this number. During the creation of IIMN networks, further layers of additional feature connections can be supplied. One example is a relationship between ion identity networks based on neutral mass differences that annotate putative structure modifications between compounds (Supplementary Figure 6). From a global view on all 24 datasets, IIMN successfully reduced the number of unconnected LC-MS^2^ features and increased the connections to annotated compound structures (Supplementary Figure 7, Supplementary Table 2).

In positive ion mode, most mass spectrometrists routinely consider H and Na adducts, but rarely NH_4_, Ca, and K adducts and in-source fragments that were commonly observed in the 24 datasets. Inspecting the relative distribution of ion identities within all datasets, marine samples, for instance, showed a higher percentage of NH_4_ adducts (24±5%) when compared to all other datasets (10±8%). Sodium adducts that were expected to be elevated in marine samples (due to anticipated higher salt contents in the original sample), in contrast, are evenly distributed between all datasets with an average of 26±6% (Figure 2g). On average, protonated species contribute to 23±6% of the overall ion identities, indicating spectral bias in public MS^2^ libraries such as MassBank of North America (66% [M+H]^+^) and GNPS (65% [M+H]^+^) (Supplementary Figure 8), and suggests that the community should provide MS^2^ spectra for other ion species of the same molecules to reference libraries. Here, IIMN can be used to expand the spectral libraries with additional adducts and in-source fragments in LC-MS experiments, which can significantly increase spectral library coverage and thus MS^2^ annotation rates. By propagating high confident spectral matches (cosine > 0.9 or authentic standards) to connected ion identities from the 24 public datasets and two datasets of natural products from the NIH ‘ACONN’ collection, we created spectral libraries with a total of 2,659 unique entries with a broader and more representative ion species coverage (e.g., 24% [M+H]+, 22% multimeric species, 17% [M+Na]^+^, 15% in-source fragments, and 13% [M+NH_4_]^+^). Such spectral libraries better represent ion species observed in typical metabolomics experiments (Supplementary Table 3 and Supplementary Figure 8).

In conclusion, by establishing relationships between different ion species originating from the same compound, IIMN facilitates molecular network interpretation and compound annotation. An exciting application of IIMN is the expansion of spectral libraries by (re)-processing public datasets and propagating spectral library annotations to create library entries of connected ion identities. The identification of ion adducts can reveal novel ionophores, some of which will be biologically relevant and are still underappreciated in the function of small molecules^12,13^. The integration into FBMN and the GNPS environment provided a platform to utilize IIMN in other related bioinformatics tools, e.g., SIRIUS^14^ and CANOPUS^15^. We anticipate that the new option to add orthogonal relationships between features to IIMN will stimulate the integration and development of additional tools for spectral alignment and measures of feature-feature relationships^16^.

To reach a broad user base, we interfaced the IIMN workflow with three widely used open source MS processing tools (MZmine^17^, MS-DIAL^18^, and XCMS^7,19^). Detailed documentation and training videos are available online (https://ccms-ucsd.github.io/GNPSDocumentation/fbmniin/). Especially the option to directly submit IIMN analysis from MZmine to GNPS provides a simple entry point for new users.

## Online Methods

### Post-column salt infusion experiments

For salt addition UHPLC-MS^2^ experiments, a mixture of 300 natural products from the NIH NCGC collection was prepared in 100 μL methanol/water/formic acid (80:19:1, Fisher Scientific, San Diego, USA) at a concentration of 0.01 μM of which 2 μL were injected into a Vanquish UHPLC system coupled to a Q-Exactive quadrupole orbitrap mass spectrometer (Thermo Fisher Scientific, Bremen, Germany) in three technical replicates. For the chromatographic separation, a reversed-phase C18 porous core-shell column (Kinetex C18, 50 x 2 mm, 1.8 um particle size, 100 Å pore size, Phenomenex, Torrance, USA) was used. For gradient elution, a Vanquish (Thermo Fisher Scientific, Bremen, Germany) high-pressure binary gradient system was used. The mobile phase consisted of solvent A H_2_O + 0.1% formic acid (FA) and solvent B acetonitrile (ACN) + 0.1% FA. The flow rate was set to 0.5 mL/min. Samples were eluted with a linear gradient from 0-0.5 min, 5% B, 0.5-8 min 5-50% B, 8-10 min 50-99% B, followed by a 2 min washout phase at 99% B and a 3 min re-equilibration phase at 5% B. Post-column we infused ammonium acetate or sodium acetate solutions (50, 5 and 0 mg/L) at 10 μL/min (dilution factor 50) with a syringe pump to yield final concentration of sodium or ammonium acetate of 1, 0.1 and 0 mg/L. Data-dependent acquisition (DDA) of MS^2^ spectra was performed in positive mode. Electrospray ionization (ESI) parameters were set to 52 psi sheath gas pressure, 14 AU auxiliary gas flow, 0 AU sweep gas flow and 400 °C auxiliary gas temperature. The spray voltage was set to 3.5 kV and the inlet capillary to 320 °C. 50 V S-lens level was applied. MS scan range was set to *m/z* 150-1500 with a resolution at *m/z* 200 of 17,500 with one micro-scan. The maximum ion injection time was set to 100 ms with an automatic gain control (AGC) target of 1E6. Up to 5 MS^2^ spectra per MS^1^ survey scan were recorded in DDA mode with a resolution of 17,500 at *m/z* 200 with one microscan. The maximum ion injection time for MS^2^ scans was set to 100 ms with an AGC target of 3.0E5 ions and a minimum 5% C-trap filling. The MS^2^ precursor isolation window was set to *m/z* 1. The normalized collision energy was set to a stepwise increase from 20 to 30 to 40% with single charge as the default charge state. MS^2^ scans were triggered at the apex of chromatographic peaks within 2 to 15 s from their first occurrence. Dynamic precursor exclusion was set to 5 s. Ions with unassigned charge states were excluded from MS^2^ acquisition as well as isotope peaks.

### Ion identity molecular networking - workflow overview

The ion identity molecular networking (IIMN) workflow aids the feature-based molecular networking workflow by adding MS^1^ specific information, which is provided as new columns in the quantification table and as additional edges in a “Supplementary Pairs” text file within the GNPS-FBMN workflow. This parameter was introduced to stimulate and facilitate the development of new computational methods that link nodes in the resulting molecular networks and was initially developed for IIMN. The text format follows a generic comma-separated style with the columns ID1 and ID2 (matching the feature IDs in the feature quantification table and mgf), EdgeType (defining the method), Score (numerical), and Annotation. To enable a broad user base to employ ion identity molecular networking in their studies, three popular mass spectrometry processing tools, namely, MZmine, MS-DIAL, and XCMS (+CAMERA), were modified with additional export scripts or modules.

### The general steps to create ion identity molecular networks

1. If needed, convert the spectral data files to an open format (e.g., mzML)
2. Import the data into one of the open-source tools: MZmine, MS-DIAL, or XCMS
3. Process the data to create a feature list (aligned over all samples)
4. Perform MS^1^-based feature grouping and ion identity annotation
5. Export the feature list as a feature quantification table (.csv), an MS^2^ spectral summary file (.mgf) which contains a representative fragmentation spectrum for each feature, and supplementary edges files (IIN files, .csv) (more information in the tool-specific workflow sections)
6. Create a metadata file to group samples for statistics (optional)
7. Upload all files to GNPS and start a new feature-based molecular networking job (MZmine can directly submit and start a new IIMN job on GNPS)
8. Download and visualize the results in a network analysis software (e.g., Cytoscape20, https://cytoscape.org/).

Refer to the documentation on how to run FBMN within GNPS and multiple mass spectrometry data processing tools.

https://ccms-ucsd.github.io/GNPSDocumentation/featurebasedmolecularnetworking/

For IIMN, refer to the related part of the GNPS documentation.

https://ccms-ucsd.github.io/GNPSDocumentation/fbmn-iin/

### IIMN with MZmine

MZmine lacked a functional algorithm to group and annotate different ion species of the same molecules. Therefore, a novel workflow was implemented and split into separate modules for feature grouping (metaCorrelate), annotation of the most common ions (ion identity networking), an option to add more ion identities to existing IINs iteratively, and modules to validate multimers and in-source fragments based on MS^2^ scans. Both the creation and expansion of ion identity networks follow customizable lists of adducts and in-source modifications to cover any type of multimers, in-source fragments, and adducts. Finally, the GNPS-FBMN export module was modified to export all needed files to run IIMN. The quant table (.csv) contains grouping and ion identity specific columns, and a new “Supplementary Pairs” text file lists all additional IIN edges. MZmine is the first tool to provide a direct submission to GNPS to start analysis jobs, consequently streamlining the workflow and lowering the entrancing energy needed to apply IIMN within GNPS.

In detail, the metaCorrelate feature grouping algorithm searches for features with similar average retention times, chromatographic intensity profiles (feature shapes) with a minimum percentage of intra-sample correlation and overlap, and minimum feature intensity correlations across all samples (Supplementary Figure 2). The feature shape correlation is a vital filter to reduce false grouping significantly and can apply either a minimum Pearson correlation (favored) or cosine similarity. A requirement is at least five data points, two on each side of the peak apex. If a low MS^1^ scan rate leads to chromatographic peaks with less than five data points, it is advisable to either redesign the acquisition method or to turn off the feature shape correlation. Note that the latter is expected to reduce the ion annotation consistency and should be used with caution. Similarly, the feature height correlation across all samples is optional, provides the same correlation or similarity measures, and additionally, relies on constant ionization conditions for all samples. Therefore, this filter should be turned off if the conditions were changed throughout the study, e.g., by changing the separation conditions or ion source parameters. The general principle of the feature height correlation is that different ions of the same molecule should follow a similar trend in abundance across all samples of the same study. If any feature, such as an [M+H]^+^ feature, increases at least 10-fold, all grouped features, e.g., [M+Na]^+^ or [M+NH_4_]^+^, should never have a negative feature height correlation coefficient and should as well increase in abundance. If both the feature shape and feature height correlation filters are omitted, feature grouping is solely filtered by the retention time window and overlap. To annotate features on an MS^1^ level, ion identity libraries are created with a user-defined list of in-source modifications (fragments and clusters), a list of adducts, and a “maximum multimers number” parameters (Supplementary Figure 2). Each adduct is combined with each modification to fill the library with ion identities for 1M to the maximum multimers number. Ion identity networks are then created by applying all ion identity pairs to all pairs of grouped features to calculate and compare the neutral masses of features *(m/z)* with specific ion identities (mass difference, charge (*z*), and multimer number). Optionally, after the creation of ion identity networks with the main library, further ion identities can be added iteratively to existing networks. This workflow enables the user to divide into commonly and uncommonly detected ion identities and ensures that each network contains at least two or more main ion identities. Finally, an ion identity network refinement provides filters for minimum network size and to only keep the largest (most descriptive) IIN per feature.

More on the integration of the new IIMN workflow in MZmine can be found online (http://mzmine.github.io/iin_fbmn).

Refer to the documentation and video tutorials on how to apply IIMN within MZmine and GNPS. The youtube playlist “MZmine: Ion Identity Molecular Networking” contains instructions on data processing for IIMN and FBMN, a minimalistic and full IIMN workflow within MZmine, and theoretical background to feature shape correlation and ion identity molecular networking. https://ccms-ucsd.github.io/GNPSDocumentation/fbmn-iin-mzmine/ https://www.youtube.com/playlist?list=PL4L2Xw5k8ITyxSyBdrcv70LDKsP8QNuyN

### IIMN with XCMS (CAMERA)

The XCMS^19^ Bioconductor package^21^ is the most widely used software for processing untargeted LC-MS based metabolomics data. Its results can be further processed with the CAMERA^7^ package to determine which of the extracted *m*/*z*-rt features might be adducts^7^ or isotopes^22^ of the same original compound. For the integration of XCMS and CAMERA into the IIMN workflow, novel utility functions were created (‘*getFeatureAnnotations’* and *‘getEdgelist’)* to extract and export MS^1^ based feature and edge annotations (i.e. grouping of features to adduct/isotope groups of the same compound). In addition, the utility function *‘formatSpectraForGNPS’* is used to export MS^2^ spectra. These functions are available in the GitHub repository https://github.com/jorainer/xcms-gnps-tools. R-markdown documents and python scripts with example analyses and descriptions are available in the documentation. (https://ccms-ucsd.github.io/GNPSDocumentation/fbmn-iin-xcms/) The files exported by these utility functions can be directly used for IIMN analysis on GNPS. Note that theoretically, it is possible to use RAMClust^8^, CliqueMS^23^, or other packages available for XCMS that perform ion annotation. The results of these packages need to be reformatted to the introduced generic supplementary edges format. The CAMERA integration might serve as a reference and starting point.

### IIMN with MS-DIAL

MS-DIAL^24^ is a polyvalent mass spectrometry data processing software capable of processing various non-targeted LC-MS metabolomics experiments, including ion mobility mass spectrometry (http://prime.psc.riken.jp/compms/msdial/main.html). MS-DIAL supports IIMN since version 4.1. After a standard data processing workflow with MS-DIAL, the “Alignment results” can be exported for IIMN analysis using the option “GNPS export”. Detailed documentation and representative tutorials are available in the GNPS documentations (https://ccms-ucsd.github.io/GNPSDocumentation/fbmn-iin-msdial).

### Dataset processing

All 24 datasets (Supplementary Table 1) were processed with the MZmine workflow. As each dataset originates from a different study and was acquired with different LC-MS methods, variable feature detection and alignment parameters were applied, which are summarized in Supplementary Table 5. For all datasets, the same parameters were used for the feature grouping module (metaCorrelate) and the ion identity networking modules, with the only exception that the feature height correlation filter was turned off to group features for the post-column salt infusion experiments. As described previously, this filter should only be applied if the ionization conditions and detection sensitivity are kept constant over all samples. The post-column infusion of different salt solutions for this study promotes the formation of specific ion species in the ionization source.

1. A pair of features were grouped with a retention time tolerance of 0.1 min, with a minimum overlapping intensity percentage of 50% in at least 2 samples in the whole dataset (gap-filled features excluded), a feature shape Pearson correlation greater equals 0.85 with at least 5 data points and 2 data points on each edge, and a feature height Pearson correlation greater equals 0.6 with at least 3 data points.
2. The initial creation of ion identity networks was performed using the ion identity networking module and a maximum tolerance of 0.001 *m/z* or 10 ppm, a comparison where a pair of features and a pair of ion identities only need to match in one sample, and an ion identity library created based on 2M as the maximum multimers number, a list of adducts ([M+H]^+^, [M+Na]^+^, [M+NH_4_]^+^, [M-H+2Na]^+^, [M+2H]^2+^, and [M+H+Na]^2+^), and a list of in-source modifications ([M-H_2_O] and [M-2H_2_O]).
3. Two iterations were applied to add more ion identities to the resulting networks of step 2 with an unchanged *m/z* tolerance.

a. To add a higher variety of adducts, a maximum multimers number of 2, a list of adducts ([M+H]^+^, [M+Na]^+^, [M+K]^+^, [M+NH_4_]^+^ [M-H+2Na]^+^, [M-H+Ca]^+^, [M-H+Fe]^+^, [M+2H]^2+^, [M+H+Na]^2+^, [M+H+NH_4_]^2+^, [M+Ca]^2+^, and [M+Fe]^2+^), and an empty list of modifications were used.
b. To add a greater variety of modifications and larger multimers, a maximum multimers number of 5, a list of adducts ([M+H]^+^, [M+NH_4_]^+^, and [M+2H]^2+^), and a list of modifications ([M-H_2_O], [M-2H_2_O], [M-3H_2_O], [M-4H_2_O], [M-HFA], and [M-ACN]) were used.

### Dataset statistics

Ion identity molecular networking statistics on all datasets were extracted with a new MZmine module and exported to a comma-separated file (csv) for evaluation in Microsoft Excel. The module is included in the special IIMN build of MZmine. All available statistics were based on the spectral input file (mgf) and the resulting network file (graphml), which was downloaded from the dataset’s corresponding GNPS results page. The graphml file contains all ion identity molecular networking results, namely, the nodes representing individual features and the edges between nodes. The mgf spectral summary file contains the corresponding MS^2^ spectrum for each feature node. While classical MN and FBMN depend on MS^2^ data for each node, IIN creates new MS^1^-based edges that might include nodes without an MS^2^ spectrum in the resulting network. For a comparison between FBMN and IIMN, only nodes present within both networks (with an MS^2^ spectrum) are considered. A statistical summary and in-depth statistics on each dataset are provided in a supplementary Microsoft Excel workbook (Supplementary File SI_IIMN_dataset_statistics.xlsx). Excerpts are summarized in Supplementary Table 2, and the different statistical measures and metadata items are described in Supplementary Table 4. One important measure is the identification density, i.e., all identified nodes and nodes with a maximum distance of n edges to at least one identified compound. Supplementary Figure 7 highlights how the additional edges of ion identity networking increase the identification density in the datasets, measured over a maximum distance of 1 to 5 edges. The increased density over one edge reflects the new links between unidentified to an identified node by IIN edge. The identification density is increased for 21 datasets, two datasets with poor identification rates exhibit no change, and one dataset lacks identifications. The maximum identification density increases over one edge of +8% results in a total of 42% of the nodes being either identified or directly linked to an identified compound. The network of the corresponding dataset, i.e., the post-column salt infusion study, contains a total of 22% identified nodes and 25% nodes with ion identity and MS^2^ spectrum in 134 ion identity networks. Ion identity molecular networking decreased the number of unconnected singleton nodes by −12% to a total of 42%. Filtering out nodes with poor MS^2^ spectra with less than four signals, which was used as the minimum number of signals for the library matching and FBMN networking, decreases the number of unconnected singleton nodes further to 29%. Consequently, the network contains many nodes without a match to any library or experimental spectra. Collapsing all nodes with IIN edges into molecular nodes reduces the total network size by −20%, which significantly reduces the overall redundancy and facilitates network visualization and analysis.

To extract the same statistics on any results from IIMN, download the networking results as a graphml file from a GNPS job page and use the mgf file of that analysis. The special MZmine IIMN build offers two modules in the tab “Tools”. More information and the latest IIMN enabled MZmine version are available (http://mzmine.github.io/iin_fbmn).

- GNPS results analysis (IIMN+FBMN)

- For a single analysis
- This tool also offers the extraction of new spectral library entries
- GNPS results analysis (IIMN+FBMN) of all sub

- For multiple analyses at once
- Generates statistics for each subfolder with exactly one graphml and mgf file (names do not have to match)

### IIMN-based spectral library generation

#### From experimental datasets

To comprehensively cover the fragmentation behavior of a molecule, spectral libraries should contain fragmentation spectra of different ion species acquired with different instrument types and fragmentation methods. IIMN might serve as a solution to expanded spectral libraries. In order to create new spectral library entries based on IIMN, all 24 datasets were searched for ion identity networks that contain a match to the GNPS spectral libraries with a minimum cosine similarity of 0.9 and a minimum number of shared fragment ions of 4-6, depending on each dataset’s FBMN parameters. For each matching IIN, all contained ion identity features with an MS^2^ spectrum and at least 3 signals above 0.1% relative intensity were extracted as new library spectra. The new library entries were constructed based on the highest library match and its attributes, namely, the compound name, structure strings as SMILES and InChI, and the neutral mass, the ion identity provided the ion species information and the precursor *m/z,* and datasetspecific metadata was added manually. With these strict rules, a total of 538 spectral entries were extracted from all 24 datasets. The new library has a broader and more distributed ion identity coverage when compared to selected representative spectral libraries from MassBank of North America (MoNA) and GNPS. At the same time, it is similar to spectral libraries that were generated with the new MSMS-Chooser library creation workflow in the GNPS ecosystem (Supplementary Fig. 5). The new IIMN-based library was made publicly available through the GNPS library batch submission (Supplementary Tab. 3).

#### From a natural product compound library

The library creation workflow was repeated and refined on the mass spectrometry data collected for the “NIH NPAC ACONN” collection of natural products (2,179 compounds) provided by Ajit Jadhav (NIH, NCATS). The IIMN workflow was optimized and then applied to two LC-MS datasets collected on mass spectrometers operating in positive ionization mode, the MSV000080492 acquired on a qTOF-MS maXis II (Bruker Daltonics, GmbH) and the MSV000083472 acquired on a Q-Exactive (ThermoFisher Scientific, MA). During feature-based molecular networking, library matching was limited to the manually created GNPS libraries, which were based on the same qTOF-MS dataset (GNPS-NIH-NATURALPRODUCTSLIBRARY, GNPS-NIH-NATURALPRODUCTSLIBRARY_ROUND2_POSITIVE, minimum matched signals=3, minimum cosine similarity=0.6). A new library for both datasets was created with new spectral entries with at least 2 signals above 0.1% relative intensity and with ion identities matching to the adduct of the library matches. Furthermore, library matches were filtered by a sample list of compound names contained in LC-MS samples. The IIMN library creation workflow resulted in 806 and 1,315 new library entries for the qTOF-MS and the Q-Exactive datasets, respectively. The new library was made publicly available through the GNPS library batch submission (Supplementary Table 3). In total, we generated 2,659 IIMN-based new spectral library entries.

#### MZmine IIMN workflow for spectral library extraction

To extract spectral library entries from any IIMN results, download the networking results as a graphml file from a GNPS job page and use the mgf file of that analysis. The special MZmine IIMN build offers the “GNPS results analysis” module in the tab “Tools” to create library entries based on these two files and provided metadata. The minimum GNPS library match score sets a threshold for the extraction of library entries. Furthermore, library matches can be filtered to also match the ion identity to the adduct of the library match. A simple comparison between the different reporting formats for adducts was implemented. It removes all spaces, square brackets, and plus symbols (e.g., harmonizing M+H and [M+H]^+^). Filters are available for new library entries with a minimum number of signals above a relative intensity threshold.

Latest information on the IIMN MS^2^ library generation workflow in MZmine is available online: http://mzmine.github.io/iin_fbmn

Documentation on the GNPS library batch submission is available at: https://ccms-ucsd.github.io/GNPSDocumentation/batchupload/

### Use case - Compound structure information

The ion identity molecular networking results for the *Stachybotrys chartarum* dataset (MSV000084134) prove that the ion identity annotations can yield structure relevant information. Putative molecular formula modifications (+O and +H_2_O) between chemical compounds can be verified by the maximum number of water losses that were annotated by IIN. The difference in the number of oxygens in the molecular formulas of phenylspirodrimane derivatives is reflected in additional losses of H_2_O within the corresponding IINs. The results are depicted in Supplementary Figure 5. The IIMN job can be accessed on GNPS (rerun of the original job after additional spectral library entries were added to the GNPS spectral libraries: https://gnps.ucsd.edu/ProteoSAFe/status.jsp?task=3bd4def5e0e348c9b113f4a072f03ea9).

### Use case - Bile acids

Networks of 88 bile acid extracts from feces and gall bladder of various animals (MSV000084170) are visualized in Supplementary Figure 4. The comparison between feature-based molecular networking with and without the additional edges from ion identity networking demonstrates how IIMN complements and improves FBMN. The new connections between different ion identities, especially between protonated and sodiated ions, merge multiple subnetworks and unconnected nodes of specific compound classes into one cluster with a higher identification density. Nodes with MS^2^ spectra that match to reference spectra of free and conjugated bile acids now fall into the same IIMN network. Finally, the complexity and redundancy are reduced by collapsing all IINs into corresponding representative nodes. The final network has a reduced number of nodes and a higher density of edges between nodes with annotations to the same compound classes. The IIMN job can be accessed on GNPS (https://gnps.ucsd.edu/ProteoSAFe/status.jsp?task=0a3f4399e5344188805e5856b756d918).

### Use case - Implementation of orthogonal supplementary edges

Ion identity molecular networking was the initial driver to implement the option of supplementary edges into the FBMN workflow on GNPS. However, based on the generic format, any tool can create and export new relationships between features to link the corresponding nodes in feature-based molecular networks. As an example, we have implemented a new MZmine module to annotate neutral mass differences between ion identity networks as putative chemical modifications, in the format of supplementary edges. These edges connect two IIN if the neutral mass difference matches a user-defined modifications list.

The IIMN MZmine workflow was applied to a dataset of 88 bile acid extracts from feces and gall bladder of various animals (MSV000084170). IIN modification edges were based on the mass differences of +methyl (Me, CH2), +O, and +H_2_O. To exemplify the results, Supplementary Figure 6 shows a network cluster of glycocholic acid analogs. Library matching annotated most of the ion identity networks as glycine conjugated bile acids; Two IINs as glycocholic acid (+isomers) and two IINs as glycodeoxycholic acid (+isomers) with a mass accuracy of <2 ppm. The additional modification edges connect these structurally related compounds and increase the network density. Moreover, they help to infer putative molecular formulas and modified structures from an FBMN. In a second analysis of the same dataset, IINs were connected based on mass differences of the modification by taurine, glycine, and alanine conjugation. This resulted, in additional links between conjugated and free bile acid forms of cholic acid and deoxycholic acid. The IIMN jobs can be accessed on GNPS.

**IIMN**

https://gnps.ucsd.edu/ProteoSAFe/status.jsp?task=0a3f4399e5344188805e5856b756d918

**IIMN: Methyl (Me, CH_2_), O, and H_2_O modification edges**

https://gnps.ucsd.edu/ProteoSAFe/status.jsp?task=465d7285380942a0828e462d1db027c2

**IIMN: Taurine, glycine, and alanine modification edges**

https://gnps.ucsd.edu/ProteoSAFe/status.jsp?task=69b40f808b2047d89fccf3d07e79fc59

### Use case - Metal-binding compounds and ionophores

Ion identity molecular networking can be used in combination with native ESI-based metabolomics^12^ to find biologically-relevant metal-binding compounds or to elucidate metalbinding preferences of known or novel metal-binding molecules (Zhi, H. et al., submitted). One recent example in which IIMN was instrumental in understanding metal-binding and selectivity is yersiniabactin. We identified yersiniabactin as a novel zincophore produced by E. coli Nissle by performing post-liquid chromatography (LC) pH adjustment (to pH 6.8) and infusion of zinc acetate solution, followed by mass spectrometry and ion identity molecular networking. With this strategy, mass spectrometry features with correlated peak shapes and retention times, in addition to an *m/z* difference resulting from zinc-binding (+Zn^2+^ -H^+^) were found. These results are summarized in Supplementary Figure 3. While this example highlights the discovery of a zinc-binding molecule explicitly, IIMN has been used in conjunction with the infusion of other metals, including iron, copper, and cobalt, to find siderophores and other ionophores. The IIMN job can be accessed on GNPS (https://gnps.ucsd.edu/ProteoSAFe/index.jsp?task=525fd9b6a9f24455a589f2371b1d9540).

### Code availability

The IIMN workflow is available as an interface on the GNPS web platform (https://gnpsquickstart.ucsd.edu/featurebasednetworking). The workflow code is open source and available on GitHub (https://github.com/CCMS-UCSD/GNPS_Workflows). It is released under the license of The Regents of the University of California and free for non-profit research (https://github.com/CCMS-UCSD/GNPS_Workflows/blob/master/LICENSE). The workflow was written in Python (ver. 3.7) and deployed with the ProteoSAFE workflow manager employed by GNPS (http://proteomics.ucsd.edu/Software/ProteoSAFe/). We also provide documentation, support, example files, and additional information on the GNPS documentation website (https://ccms-ucsd.github.io/GNPSDocumentation/), and we invite everyone to contribute to the documentation on GitHub.

The source code of all modules which were implemented into MZmine, e.g., the Export for IIMN module, the metaCorrelate grouping module, the ion identity networking modules, and the results and spectral library generation module, is available at http://mzmine.github.io/iin_fbmn under the GNU General Public License. The source code for the custom GNPS export functions for XCMS is available at https://github.com/jorainer/xcmsgnps-tools under the GNU General Public License.

### Data availability

All raw (.raw) and peak picked (.mzXML or .mzML) mass spectrometry data as well as processed data (.mgf and .csv) and ion identity molecular networks are available through the MassIVE repository (massive.ucsd.edu). Individual MassIVE dataset identifiers are listed in Supplementary Table 1. Dataset metadata and MZmine processing parameters are available in Supplementary Table 5. The statistical results on all 24 datasets are available in Supplementary File SI_IIMN_dataset_statistics.xlsx. The ion identity statistics on different MS^2^ spectral databases are available as Supplementary File SI_IIMN_spectral_library_anaylsis.xlsx.

## Acknowledgments

We thank the German Chemical Industry Fund (FCI, Fonds der Chemischen Industrie) for a Ph.D. scholarship and travel support to RS. We thank the Deutsche Forschungsgemeinschaft for support to DP (PE 2600/1-1) and to SB and KD (BO 1910/20). AB thanks FAPESP fellowship (2018/24865-4). COG thanks FAPESP scholarship (2019/06061-8). PCD was supported by the Gordon and Betty Moore Foundation (GBMF7622), the US National Institutes of Health for the Center (P41 GM103484, R03 CA211211, R01 GM107550). LFN was supported by the US National Institutes of Health (R01 GM107550), and the European Union’s Horizon 2020 program (MSCA - GF, 704786). AMCR and PCD were supported by the National Sciences Foundation grant IOS-1656481. LIA was supported by the National Science Foundation grant OCE-1736656. MR is supported by Public Health Service Grants AI126277, AI114625, AI145325, by the Chiba University-UCSD Center for Mucosal Immunology, Allergy, and Vaccines and an Investigator in the Pathogenesis of Infectious Disease Award from the Burroughs Wellcome Fund. DP, MAP and KIP were supported by the National Science Foundation’s Center for Aerosol Impacts on the Chemistry of the Environment (CAICE) under grant number CHE1801971. KLM was supported by the Gordon and Betty Moore Foundation (GBMF6920) and the US National Institutes of Health (R01 GM132649). RT was supported by the US National Institutes of Health (NCCIH T32AT010131). AAO acknowledges the support of Fulbright Commission and Consejo Nacional de Investigaciones Científicas y Técnicas (CONICET-Argentina). ZK was supported by the program Lumina Quaeruntur of the Czech Acad Sci. ALG was supported by Vaincre la mucoviscidose and Association Grégory Lemarchal. HMR thanks CNPq (#142014/2018-4), and the Brazilian Fulbright Commission for the scholarships provided. We thank Ajit Jadhav (NIH/NCATS) for providing the compounds used for the adduct induction experiment, and for the library generation. We thank Andreas J Andersson, Heather N Page, Travis A Courtney, Evan Fox, Sara P. Pucket, Kathleen E. Kyle, Jonathan L. Klassen and Marcy J. Balunas, Andrea Fidgett and Michelle Gaffney for providing samples and assisting during sampling campaigns.

