## Supplementary Figure for "Ion Identity Molecular Networking in the GNPS Environment"

### Content:

Page 3-9: Supplemental Figures
Page 9-18: Supplemental Tables
Page 19-20: Supplemental References

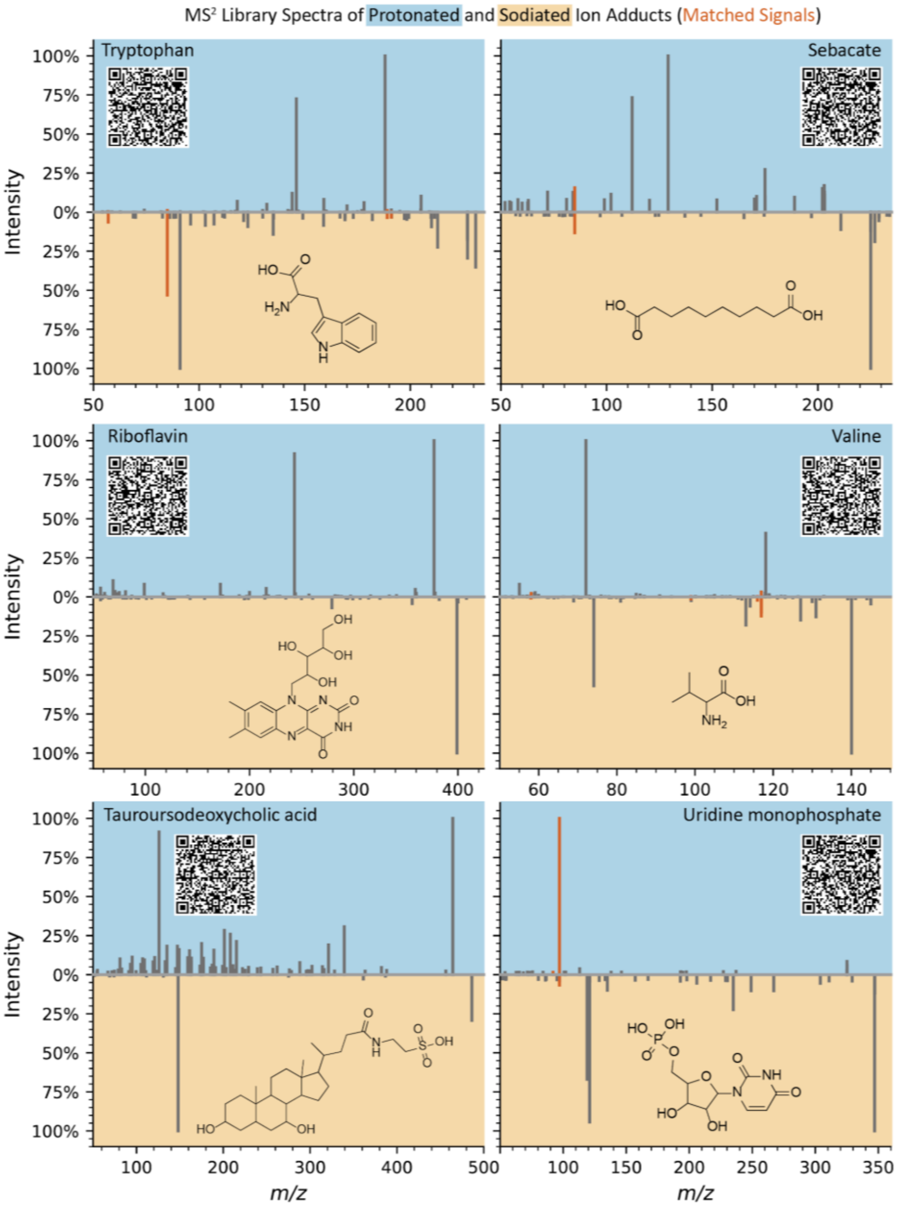


**Supplementary Figure 1: Comparison of MS$^2$ library spectra for sodiated and protonated ion species of six compounds.** The spectral mirror charts show MS$^2$ library spectra of the same compound as $[M+H]^+$ (top plot) and $[M+Na]^+$ (bottom plot). Due to high in-source fragmentation and low $[M+H]^+$ abundance, tauroursodeoxycholic acid compares the $[M-H_2O+H]^+$ and $[M+Na]^+$ MS$^2$ library spectra. Matched signals in mirror charts are highlighted (dark orange). These charts exemplify a commonly seen difference in fragmentation behavior for sodiated and protonated ion adducts. All library spectra are publicly available in the GNPS spectral libraries. The charts were created with the Metabolomics Spectrum Resolver (https://metabolomics-usi.ucsd.edu/) and scannable QR-codes link out to the individual annotated charts and library spectra with their metadata.

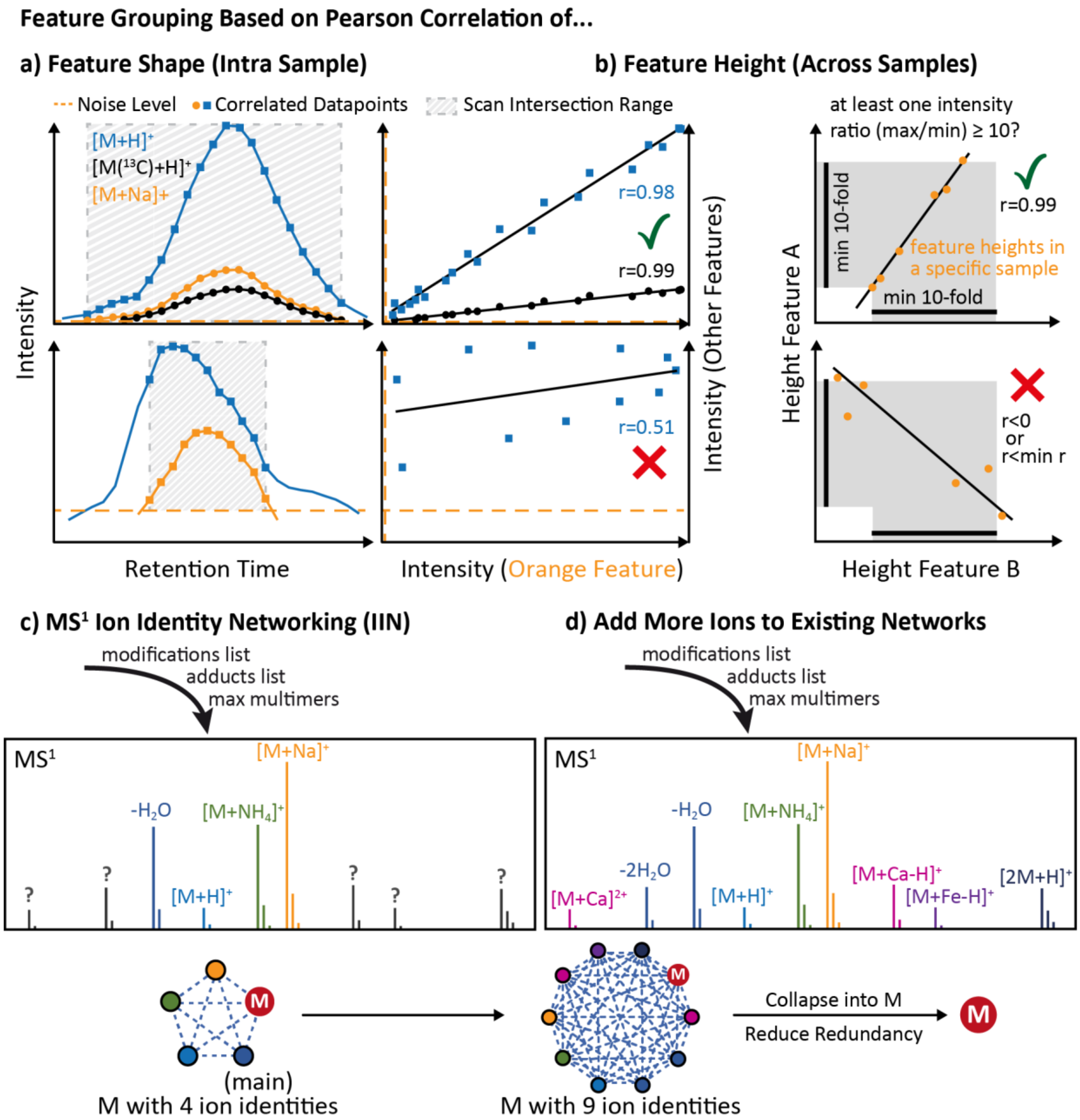


**Supplementary Figure 2: Schematic workflow for feature grouping (metaCorrelate) and ion identity networking within MZmine.** After preselecting features that are in a retention time window and have a minimum overlapping retention time percentage (measured on the smaller feature for each tested pair), features are grouped if **a)** their shapes (intensity profiles) and **b)** their heights across all samples resolve to Pearson correlation coefficients (r) of user-defined thresholds. Optionally, both correlation filters can be omitted or switched to a cosine score similarity threshold. By turning off both filters, the grouping is solely based on the retention time window and overlap prefilters. Ion identity networking uses an in-source modifications list, an adducts list, and a “maximum multimers number” parameter to create a library of $MS^1$ ion identities. To create annotation networks, all pairs of ion identities are applied to all pairs of grouped features to calculate the neutral mass of the corresponding feature ($m/z$) with an ion identity (mass difference, charge ($z$), and multimer number). After **c)**, the initial creation of ion identity networks with a user-defined library of main ion identities, more uncommon ion identities can **d)** be added iteratively to existing networks.

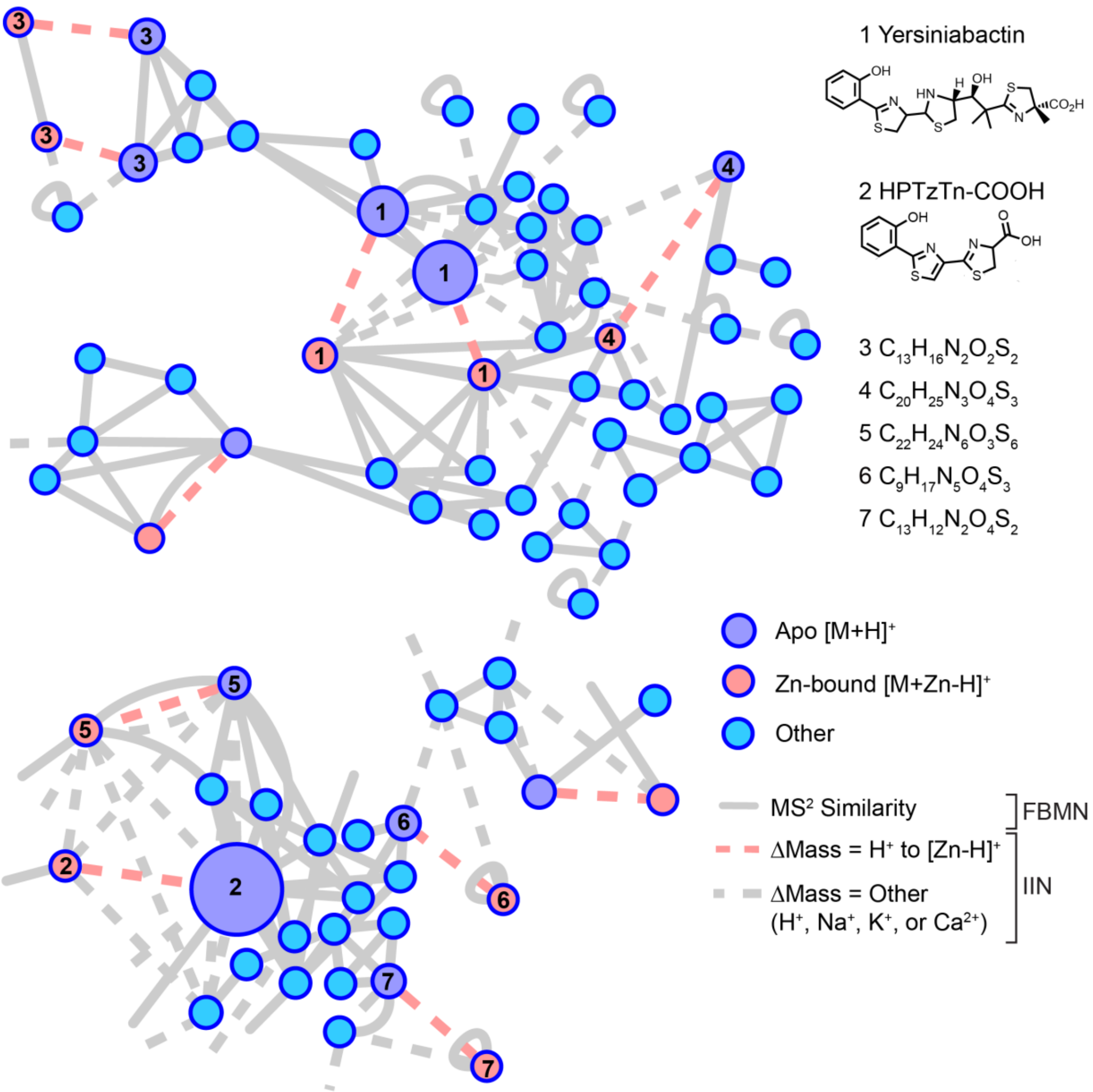


**Supplementary Figure 3: Use case for the discovery of metal-binding compounds and ionophores.** IIMN in conjunction with native spray metal metabolomics[12] facilitated the discovery of yersiniabactin as a zincophore produced by *E. coli* Nissle (Zhi, H. et al., *submitted*). Zinc-binding molecules, such as yersiniabactin and other potential derivatives or truncations, are shown in salmon, while the corresponding protonated form of these molecules is shown in purple; other nodes are colored light blue. Structures and molecular formulas (generated using SIRIUS 4.0) for compounds 1-7 are provided.

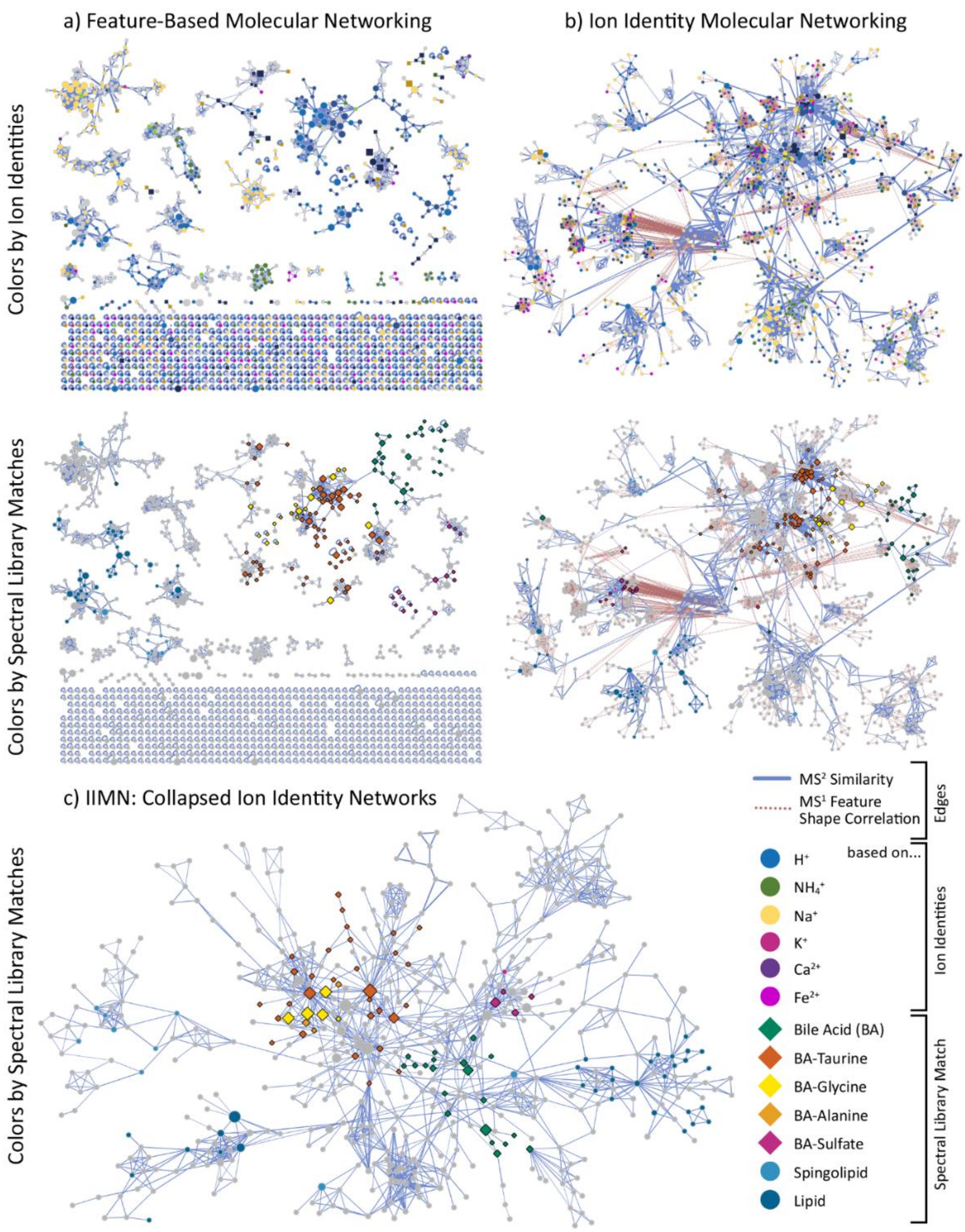


**Supplementary Figure 4: Network comparisons for a cluster with matches to bile acids from 88 bile acid extracts from feces and gall bladder of various animals (MSV000084170).** This overview compares the feature-based molecular networking (FBMN) results a) without and b) with ion identity networking (IIN) and after c) collapsing all ion identity networks into single representative nodes. In the upper two networks, nodes are colorized depending on which adduct the detected ion identities are based on. In contrast, the lower three networks emphasize nodes with $MS^2$ spectra that match specific compound classes, mainly bile acids and different conjugates. These results prove that FBMN is a suitable method to connect structurally similar compounds, such as isomers, based on $MS^2$ spectral similarity scoring. However, only one edge was established between clusters of free and conjugated bile acids. Overall, bile acid analogs were separated into multiple subnetworks and unconnected nodes with a clear trend of separation into clusters of sodiated and protonated ion identities. Ion identity molecular networking added new edges between ion identities of the same molecules. This results in a higher network density with more connections between annotated and unknown nodes, fewer unconnected nodes, and clustering of compound classes. The collapsed network c) reduces the complexity and redundancy of having multiple nodes per compound and only keeps $MS^2$ spectral similarity edges.

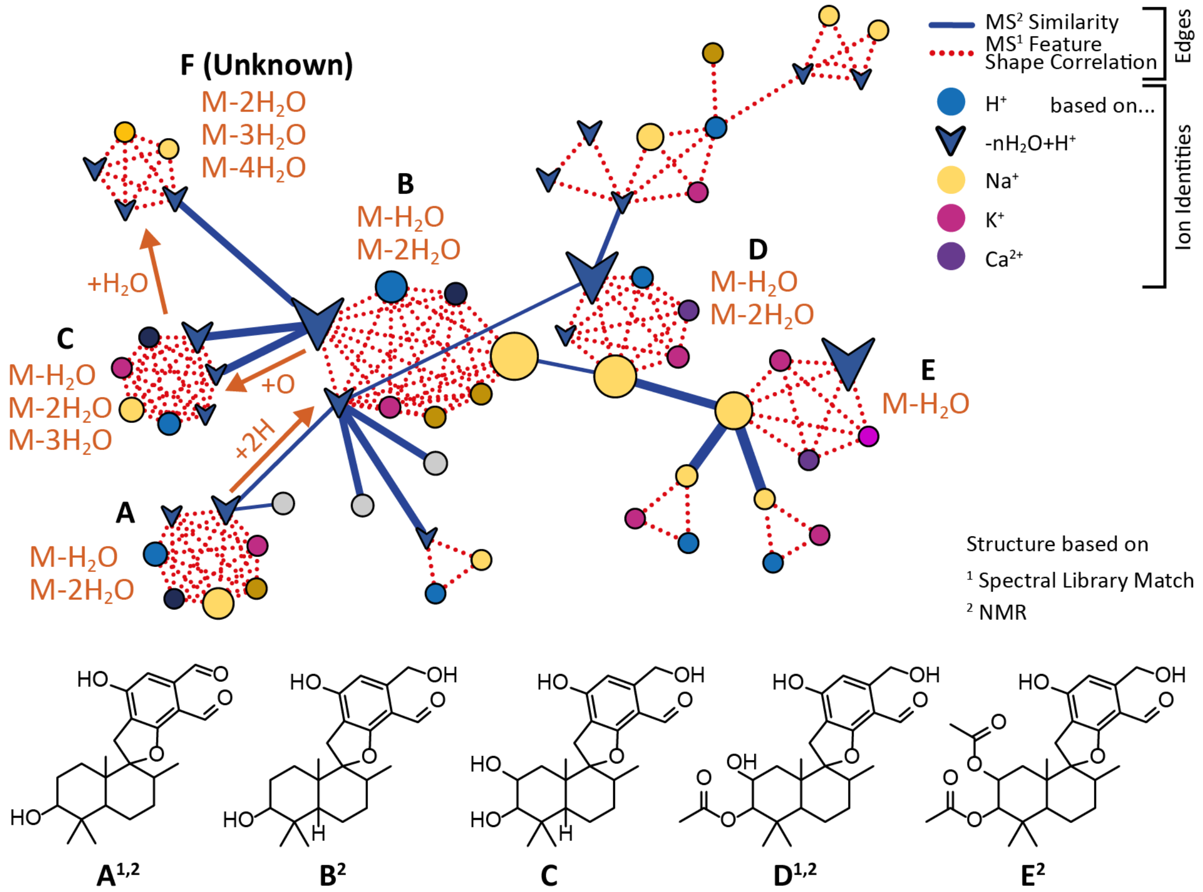


**Supplementary Figure 5: Ion identity molecular networking results for *Stachybotrys chartarum* liquid culture extracts (MSV000084134).** Compounds A and D were annotated by spectral library match and compounds A, B, D, and E were verified by nuclear magnetic resonance (NMR) spectroscopy[27]. Modifications between compounds are based on the compound structures or the differences of the average neutral masses for each IIN. The modifications of +O and $+H_2O$ from B to C to F can also be deduced from the ion identity annotations, as the maximum in-source water losses for each compound confirm the addition of oxygens; Compounds A, B, and D ($-2H_2O$ maximum water losses), compound C ($-3H_2O$), and compound F ($-4H_2O$). Therefore, this example proves that IIN can yield structure relevant information and facilitate structure annotation.

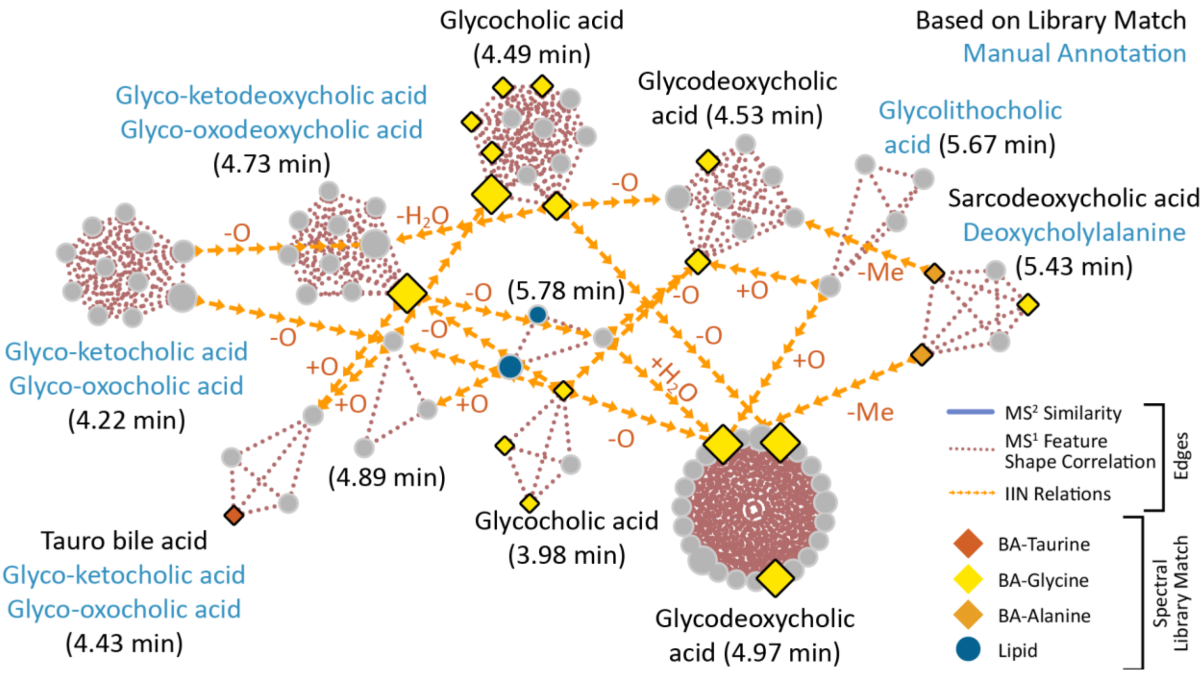


**Supplementary Figure 6: Use case for supplementary edges to spot compound modifications.** Ion identity networks were linked by additional edges based on neutral mass differences matching those of methyl (Me, $CH_2$), O, or $H_2O$. $MS^2$ spectral similarity edges are hidden to reduce visualization complexity. Spectral library matching annotated two IINs as glycocholic acid (+isomers) and two IINs as glycodeoxycholic acid (+isomers). The new additional modification edges helped to infer putative annotations for six IINs, moreover, in the resulting IIMN, related compounds clustered closer and with more connections.

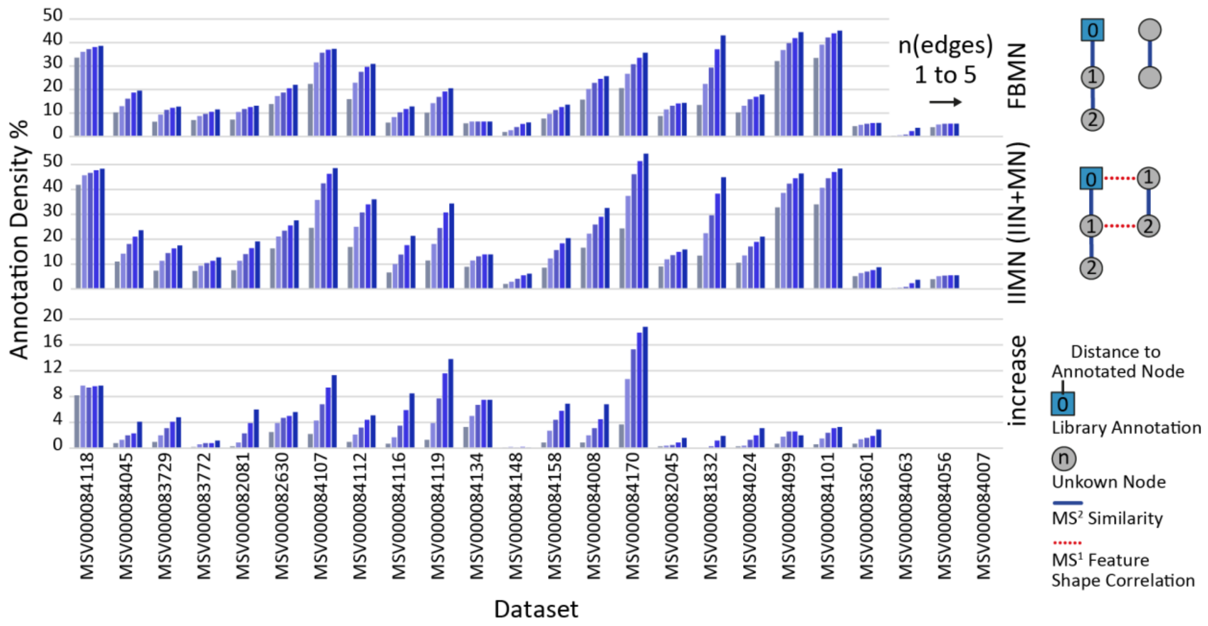


**Supplementary Figure 7: Compound annotation densities for maximum distances over 1 to 5 edges to an annotated node.** From top to bottom, nodes that are connected to at least one annotated compound by n FBMN edges (top), by IIMN (IIN+FBMN) edges (middle), and the differential increase of annotation density by adding IIN to FBMN (bottom). The increase over one edge corresponds to unknown features (nodes) which are directly connected by an IIN edge to an annotated compound (maximum increase +8%).

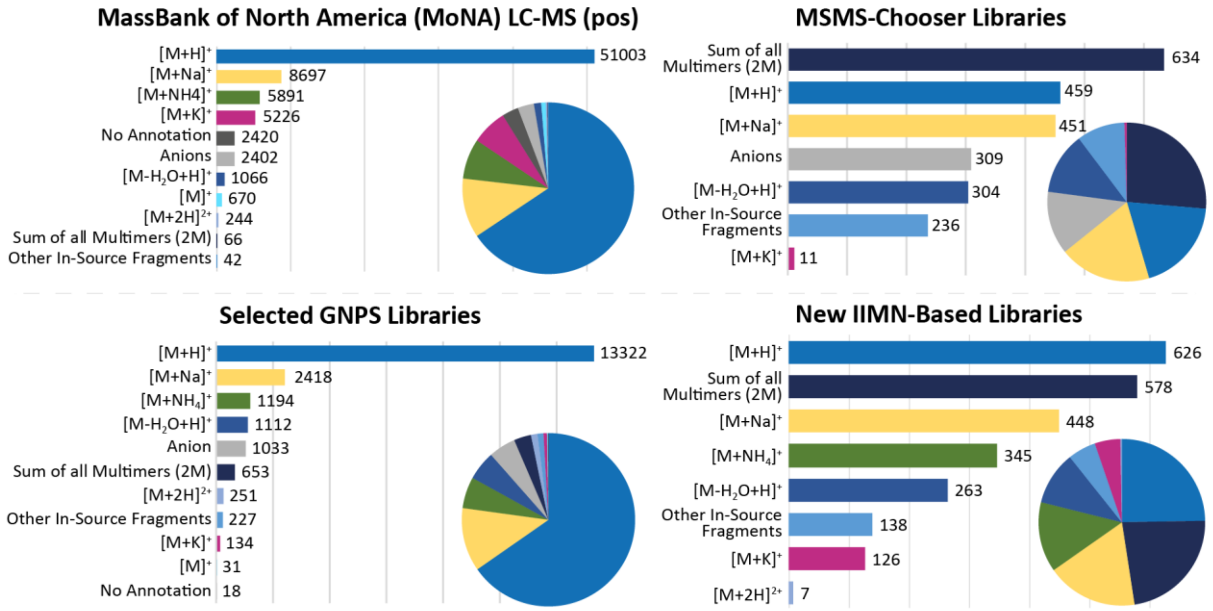


**Supplementary Figure 8: Analysis of the coverage and distribution of ion identities in public LC-MS$^2$ spectral libraries** (refer to Supplementary Table 3 for library origins). Two-thirds of the MassBank of North America LC-MS$^2$ positive ion mode library entries were entered as $[M+H]^+$ while only four other ion types reach more than 1000 entries, namely, $[M+Na]^+$, $[M+NH_4]^+$, $[M+K]^+$, and $[M-H_2O+H]^+$. Other in-source fragments, multiply-charged species, and multimers are only covered for a few compounds. A significant number of entries were either annotated as negatively charged adducts (e.g., M-H) or were missing an annotation. As the ion identity naming was not harmonized, different versions pointing to the same ion identity were added to a total count. A similar ion annotation coverage was found in the GNPS spectral libraries. In contrast, libraries that were generated with the recently described MSMS-Chooser[26] workflow on GNPS or the here described IIMN library extraction workflow show an overall broader coverage of different adducts, multimers, and in-source fragments. The depicted statistical visualization compares a subset of significant or representative ion identities. The IIMN-based numbers summarize the libraries from both the 24 experimental datasets and the two NIH natural product standards datasets with a total of 2659 library entries.

**Supplementary Table 1**: LC-MS$^2$ datasets used for the evaluation of ion identity molecular networking. All datasets are publicly available through their corresponding ID in the MassIVE repository (massive.ucsd.edu).

| # | Dataset Name | Sample Origin | Dataset Description | Organism Name | MassIVE ID |
|---|---|---|---|---|---|
| 1 | B. Sub knockout | *B. subtilis* | Extracts from B. sub wildtype and srf-KO cell culture | *Bacillus subtilis* | MSV000082081 |
| 2 | Coral Reef Sea Water | Seawater | Spatial survey of Coral Reef in Hawaii, filtered and unfiltered samples | Seawater | MSV000084116 |
| 3 | Drug Metabolism Tsunoda | Human | Human blood (plasma) prepared using Phree kit, individuals given 4 different drugs (omeprazole, dextromethorphan, caffeine, and midazolam) | *Homo sapien* | MSV000084008 |
| 4 | SD Cheetahs | Cheetah | Fecal samples from cheetahs in human care | *Acinonyx jubatus* | MSV000084099 |
| 5 | Nascent Sea Spray Aerosol | Seawater | isolated sea spray aerosol | Seawater | MSV000084119 |
| 6 | Fungus-Growing T. septentrionalis Gardens | Ants fungus garden extracts | DCM:MeOH 2:1 extracts from Trachymyrmex septentrionalis fungus gardens. | *Trachymyrmex septentrionalis* | MSV000084024 |
| 7 | HMP cultures full set | Various bacterial cultures | Extracts of HMP project isolates that were grown successfully in three different media | 433 various bacteria | MSV000082045 |
| 8 | Foodomics subset | Diverse foods | Subset of food samples from Global FoodOmics project | Various foods | MSV000084101 |
| 9 | Stromatolites tissue | Intertidal - freshwater/marine stromatolites | Extracts from South African stromatolite cores, and San Diego | Cyanobacteria/Heterotrophic bacteria | MSV000083729 |
| 10 | Stromatolites DOM | Intertidal - freshwater/marine DOM | DOM/TOM analysis of water from SA stromatolite barrage pool | Cyanobacteria/Heterotrophic bacteria | MSV000083772 |
| 11 | Bacteria from Rocas Atoll | Sediments from Rocas Atoll | Extracts produced by marine bacteria in liquid cultures | Bacteria | MSV000083601 |
| 12 | B. subtilis extracted cell-free supernants | *B. subtilis* | Methanol extracts from cell-free supernatants of B. subtilis wild strains | *Bacillus subtilis* | MSV000084045 |
| 13 | SEED Grant - Sejal - Urine | Human | Urine samples extracted in 80% MeOH of Disease-associated microbial species, for disease diagnosis and prevention of Type I diabetes | *Homo sapien* | MSV000084112 |

**Supplementary Table 1: continued**

| # | Dataset Name | Sample Origin | Dataset Description | Organism Name | MassIVE ID |
|---|---|---|---|---|---|
| 14 | ONR Primary Wright - Human-Plasma | Human | Dataset investigating the impact of sleep and circadian disruption on human plasma | *Homo sapien* | MSV000082630 |
| 15 | Basidiobolus | Fungi, Zoopagomycota | MeOH extract/RP-SPE fractions of B. meristosporus, grown in 7 growth conditions in triplicate. EtOAc extract/RP-SPE fractions of culture broths. Study aims to discover the product of bacterial-like NRPSs. | *Basidiobolus meristosporus* | MSV000084007 |
| 16 | Pseudonocardia Extracts Grouping - Illumina | *Pseudonocardia* | EtOAc extracts of 7 Pseudonocardia strains to discover if there some kind of grouping according to metabolomic content | *Pseudonocardia* | MSV000084056 |
| 17 | R_HAN_01.05 | *Mus musculus* | Plasma serum from mice fed with prebiotics or antibiotics for 3 weeks | *Mus musculus* | MSV000084107 |
| 18 | Tunicates - Euherdmania X Eudistoma vannamei | Marine organisms | Extraction with MeOH of macerated organism | *Euherdmania* and *Eudistoma vannamei* | MSV000084063 |
| 19 | Saliva samples from caries and healthy children | Human | Saliva was lyophilized and non-polar molecules were extracted in ethyl acetate | Oral microbiome | MSV000081832 |
| 20 | Stachybotrys-WT-OE | Filamentous fungi | Extracts from mycelium and filtates | *Stachybotrys chartarum* | MSV000084134 |
| 21 | Urine biomarker discovery | Human | Diluted urine | *Homo sapiens* | MSV000084148 |
| 22 | SeaScape2019 Bloom1+2 | Seawater | Non-targeted metabolomics (DOM, 02 um surpore) from bulk water samples (1L) from SeaScape2019 Bloom1+2 from PPL(200mg) solid phase extraction. [doi:10.25345/C5HH23] | Seawater | MSV000084158 |
| 23 | Animal bile acids | Animals (Lee Hagey collection) | Methanolic extracts of gall bladder or feces | Various *vertebrata* | MSV000084170 |
| 24 | Adduct induction | Natural Products | Post-column salt addition | Standards | MSV000084118 |

**Supplementary Table 2:** Summary of statistical results on all 24 datasets. Refer to the provided Microsoft Excel workbook (SI_IIMN_dataset_statistics.xlsx) for a full dataset resolved statistical summary and in-depth results for each dataset.

| **Statistical measure** | **Mean %** | **Median %** | **Min %** | **Max %** |
|---|---|---|---|---|
| Library matches | 6 | 4 | 0 | 23 |
| Ion identities | 14 | 14 | 2 | 43 |
| Ion identities with $MS^2$ | 12 | 11 | 2 | 35 |
| Library matches with ion identity | 16 | 16 | 0 | 75 |
| Singletons (only FBMN edges) | 43 | 39 | 23 | 91 |
| Singletons (all edges) | 38 | 34 | 22 | 80 |
| Reduced singletons | 5 | 3 | 0 | 30 |
| Nodes reduced by IIN | 9 | 8 | 1 | 23 |

**Supplementary Table 3: LC-$MS^2$ spectral library summary.** For the statistical analysis of the ion identity coverage in available online resources, different spectral libraries were downloaded from MassBank of North America (MoNA) and GNPS and compared to the IIMN-based library, from the described workflow. MSMS-Chooser is a new workflow in the GNPS environment that focuses on the creation of spectral libraries with broader coverage of different ion identities. The currently available libraries based on MSMS-Chooser mainly contain spectral $MS^2$ entries for bile acids. All resources were downloaded on Dec, 17th 2019.

| Library Group | Library Name | Link |
|---|---|---|
| MoNA[25] | LC-$MS^2$ (positive) | https://mona.fiehnlab.ucdavis.edu/downloads |
| GNPS[2] | • GNPS Library<br>• FDA Library Pt 1<br>• FDA Library Pt 2<br>• PhytoChemical Library<br>• NIH Clinical Collection 1<br>• NIH Clinical Collection 2<br>• NIH Natural Products Library Round 1 (NIH NCGC)<br>• NIH Natural Products Library Round 2 (NIH NPAC ACONN)<br>• Pharmacologically Active Compounds in the NIH Small Molecule Repository<br>• Faulkner Legacy Library provided by Sirenas MD<br>• EMBL Metabolomics Core Facility (EMBL MCF)<br>• Pesticides<br>• Medicines for Malaria Venture Pathogen Box<br>• LDB Lichen Database<br>• GNPS Collections Miscellaneous<br>• GNPS Collections Bile Acid Library 2019<br>• MIADB Spectral Library | https://gnps.ucsd.edu/ProteoSAFe/libraries.jsp |
| MSMS-Chooser-based (GNPS)[26] | • GNPS-MSMLS<br>• GNPS Collections Bile Acid Library 2019 | https://gnps.ucsd.edu/ProteoSAFe/gnpslibrary.jsp?library=GNPS-MSMLS<br>https://gnps.ucsd.edu/ProteoSAFe/gnpslibrary.jsp?library=BILELIB19 |
| IIMN-based | • Ion identity molecular networking-based library extracted from 24 publicly available datasets<br>• Ion identity molecular networking-based library generated from the NIH Natural Products Library (NIH NPAC ACONN). All spectral entries were added to the existing GNPS library (https://gnps.ucsd.edu/ProteoSAFe/gnpslibrary.jsp?library=GNPS-NIH-NATURALPRODUCTSLIBRARY_ROUND2_POSITIVE). | https://gnps.ucsd.edu/ProteoSAFe/status.jsp?task=463118b73bad48518f124d09a86787de<br>https://gnps.ucsd.edu/ProteoSAFe/status.jsp?task=904e6d42b5024c5cacef6dd86f02b714<br>https://gnps.ucsd.edu/ProteoSAFe/status.jsp?task=c39d788d30f9408a9e53e68ec84868c6 |

**Supplementary Table 4:** Description of all statistical measures that were extracted for each dataset.

| Measure | Description |
|---|---|
| Metadata | |
| MassIVE ID | ID to access the publicly available dataset in the MassIVE repository |
| Sample origin | Study and sample descriptor (e.g., Standards, Urine,..) |
| MS type | Mass analyzer type (FTMS or qTOF) |
| Machine type | Name of the mass spectrometer |
| Summary of IIMN+FBMN statistics | |
| Total nodes | All feature nodes (might include nodes without $MS^2$) |
| Total nodes with $MS^2$ | All feature nodes with $MS^2$ spectrum |
| Library matches ($MS^2$) | Matches to the GNPS spectral libraries (= annotations) |
| Ion identities ($MS^1$) | Nodes with ion identity annotation based on $MS^1$ |
| Ion identities with $MS^2$ | Nodes with ion identity annotation that have an $MS^2$ spectrum |
| $MS^2$ with ion identity % | Nodes with ion identity relative to total nodes with $MS^2$ |
| Library matches + ion identity | Library matches with ion identities |
| Library matches + ion identity % | Relative to all library matches |
| Ion identity networks | Number of ion identity networks, representing connected ion identities which point to the same neutral molecular mass |
| All measures below were calculated in regards to nodes with a minimum number of $MS^2$ signals, filtered with n=0,1,4,6 signals. FBMN and IIMN comparisons are based on results for n=0 (no filter). | |
| $MS^2$ scans | Nodes with a spectrum with at least n signals |
| Annotated | Annotated nodes with at least n $MS^2$ signals |
| Annotated % | Relative to all nodes with $MS^2$ scans with at least n signals |
| Singletons FBMN / IIMN | Nodes that have no connecting FBMN edge / nodes that have no connecting IIMN edge |
| Singletons FBMN / IIMN% | Relative to all nodes with $MS^2$ scans with at least n signals |
| Nodes reduced by IIN | Redundant nodes that describe the same neutral molecule are reduced by collapsing all ion identities of an IIN into a singular neutral molecule node “M”. An IIN with 3 ions (nodes) would then result in a reduction of 2 nodes for the remaining “M”-node. |
| Nodes reduced by IIN % | Relative to all nodes with $MS^2$ scans with at least n signals |

**Supplementary Table 4: continued**

| Measure | Description |
|---|---|
| Remaining nodes after reduction | $MS^2$ scans - nodes reduced by IIN |
| Possible new library spectra by IIN | Counts all nodes ($MS^2$ spectra) without a direct match to the GNPS spectral libraries but with an IIN edge to at least one identified node. The count will be 2 if one or more nodes in an ion identity network are identified by a spectral library match and connected to 2 additional unidentified ion identities with at least n $MS^2$ signals. |
| Identification density as the distance to an annotated node (each node only counts once even if connected to multiple annotated compounds) in regards to nodes with a minimum number of $MS^2$ signals, filtered with n=0,1,4,6 signals. FBMN and IIMN comparisons are based on results for n=0 (no filter). | |
| Distance 0,1,2,3,4,5 edges | Number of nodes with a minimum distance (0-5 edges) to an identified node with a spectral library match. Comparison between FBMN $MS^2$-based edges only and IIMN ($MS^2$+$MS^1$-based) networks. |
| Distance 0,1,2,3,4,5 edges % | Relative to all nodes with $MS^2$ scans with at least n signals (identification density over n edges) |
| Ion identity networks statistics | |
| Adduct distribution | List of all ion identities and their occurrence in the dataset |
| NetID | The network identifier number. All nodes with an ion identity point to a specific netID. |
| Net size | Number of feature nodes in a network |
| Nodes with $MS^2$ | Number of nodes with an $MS^2$ spectrum |
| Identified | Nodes with a match to the GNPS spectral libraries |
| All ions | All ion identities (e.g., $[M+H]^+$) |
| Ions matched library | All ion identities with a library match |
| Reduction by | Reduction of nodes by collapsing IINs into a single neutral molecule-node. (Nodes with $MS^2$-1) |
| Possible new library entries with n signals (n=3,4,6) | If IIN has at least one identified node: List of ion identities with no match to the GNPS spectral libraries but at least n $MS^2$ signals. |

**Supplementary Table 5**: MZmine parameters for LC-$MS^2$ dataset processing.

| ID | Samples | MS Instrument | MS Resolution | Gradient [min] | Duty Cycle Time | $MS^1$ threshold | $MS^2$ Threshold | MS Tolerance | RT Tolerance [min] | Chromatogram Deconvolution Algorithm | $MS^2$ Paring *m/z* Tolerance | $MS^2$ Paring RT Tolerance [min] | Feature Aligment (m/z Tolerance, RT Tolerance, m/z : RT Weights) | Duplicate Row Filter | Min. Peaks in a Row | Min. Isotope Pattern Peaks | $MS^2$ Filtered | Gap Filled |
|---|---|---|---|---|---|---|---|---|---|---|---|---|---|---|---|---|---|---|
| MSV000082081 | 7 | Q-Exactive | 17000 | 5 | <1sec | 1E5 | 1E3 | 10 ppm | 0.02 | Baseline cut-off (Min peak height: 3E5, Peak dur 0.01 - 3 min, Baseline level 1E5) | 0.01 | 0.1 | 10 ppm, 0.2 min, 75:25 | no | 2 | 2 | MS2 filter | yes |
| MSV000084116 | 64 | Q-Exactive | 17000 | 15 | <1sec | 1E5 | 1E3 | 10 ppm | 0.02 | Baseline cut-off (Min peak height: 3E5, Peak dur 0.01 - 3 min, Baseline level 1E5) | 0.02 | 0.1 | 10 ppm, 0.15 min, 75:25 | no | 2 | 2 | no | yes |
| MSV000084008 | 439 | QToF/ Maxis Impact HD | | 7.5 | | 1E3 | 1E2 | 0.01 m/z or 20 ppm | 0.05 | Local minimum search (5.0%, 0.05 min, 5.0%, 3000, 2, 0.05 - 2.00 min) | 0.025 | 0.01 | 0.01 m/z or 20.0 ppm, 0.1 min, 50:25 | yes | 2 | 2 | only with MS2 or annotation | yes |
| MSV000084099 | 556 | QToF/ Maxis Impact HD | | 12.5 | | 2E3 | 9E1 | 0.001 m/z or 20 ppm | 0.03 | Local minimum search (96%, 0.03 min, 5.0%, 2000, 1, 0.0 - 2.00 min) | 0.02 | 0.15 | 0.0015 m/z or 15 ppm, 0.2 min, 2:1 | yes | 2 | 2 | no | no |
| MSV000084119 | 26 | Q-Exactive | 70000 | 10 | <1s | 1E5 | 1E3 | 5 ppm | 0.1 | local minimum search (1%, 0.05min, 1%, 1E5, 1, 0.1 - 3.00min) | 0.01 | 0.2 | 10 ppm, 0.2 min, 75:25 | no | 2 | 2 | MS2 filter | yes |
| MSV000084024 | 251 | QToF/ Maxis Impact | | 8 | 3Hz MS1/ 10Hz MS2 | 1E4 | 1E2 | 25 ppm | 0.01 | Baseline cut-off (Min peak height 1.0E4; peak duration range 0.01-1.0 min; Baseline level 1.0E2) | 0.01 | 0.3 | 25 ppm, 0.2 min, 75:25 | no | 2 | 2 | only with MS2 or annotation | no |
| MSV000082045 | 2249 | QToF/ Maxis Impact | 17000 | 10 | <1s | 1E3 | 1E2 | 20 ppm | 0.05 | Local minimum search (5.0%, 0.05 min, 5.0%, 3000, 2, 0.05 - 2.00 min) | 0.01 | 0.05 | 20 ppm, 0.1 min, 50:34 | yes | 2 | 2 | MS2 filter | yes |
| MSV000084101 | 411 | QToF/ Maxis Impact | | | | 2E3 | 9E1 | 0.001 m/z or 20 ppm | 0.03 | Local minimum search (96%, 0.03 min, 5.0%, 2000, 1, 0.0 - 2.00 min) | 0.02 | 0.15 | 0.0015 m/z or 15ppm, 0.2 min, 2:1 | no | 2 | 2 | no | no |
| MSV000083729 | 298 | Q-Exactive | 17000 | | <1s | 1E5 | 1E3 | 0.005 m/z or 10 ppm | 0.02 | Local minimum search (1%, 0.05min, 1%, 3000, 1.5, 0.05 - 2.00min) | 0.01 | 0.1 | 0.01 m/z or 10ppm, 0.2 min, 75:25 | no | 2 | 2 | no | no |
| MSV000083772 | 80 | Q-Exactive | 17000 | | <1s | 1E5 | 1E3 | 0.005 m/z or 10 ppm | 0.02 | Local minimum search (1%, 0.05min, 1%, 3000, 1.5, 0.05 - 2.00min) | 0.01 | 0.1 | 0.01 m/z or 10ppm, 0.2 min, 75:25 | no | 2 | 2 | no | no |

176 **Supplementary Table 5: continued**

| ID | Samples | MS Instrument | MS Resolution | Gradient [min] | Duty Cycle Time | $MS^1$ threshold | $MS^2$ Threshold | MS Tolerance | RT Tolerance [min] | Chromatogram Deconvolution Algorithm | $MS^2$ Paring *m/z* Tolerance | $MS^2$ Paring RT Tolerance [min] | Feature Aligment (m/z Tolerance, RT Tolerance, m/z : RT Weights) | Duplicate Row Filter | Min. Peaks in a Row | Min. Isotope Pattern Peaks | $MS^2$ Filtered | Gap Filled |
|---|---|---|---|---|---|---|---|---|---|---|---|---|---|---|---|---|---|---|
| MSV000083601 | 76 | Q-ToF/micrO TOF-QII | 18000 | 25 | | 1E3 | 1E2 | 20 ppm | 0.02 | Local minimum search (1.0%, 0.15 min, 1.0%, 1000, 1.5, 0.10 - 2.00 min) | 0.05 | 0.2 | 20 ppm, 0.3 min, 75:25 | no | 2 | | NA | no |
| MSV000084045 | 11 | Q-Exactive | 17000 | 5 | <1s | 1E5 | 1E3 | 10 ppm | 0.02 | Baseline cut-off (Min peak height: 1E5, Peak dur 0.01 - 2 min, Baseline level 1E5) | 0.01 | 0.1 | 0.01 m/z or 10 ppm, 0.1 min, 75:25 | no | 2 | 2 | only with MS2 or annotation | yes |
| MSV000084112 | 116 | Q-Exactive | 32000 | 12.5 | | 1E5 | 5E2 | 0.01 m/z or 20 ppm | 0.05 | Local minimum search (0.01%, 0.04 min, 0.01%, 1E5, 2 0.05 - 0.50 min) | 0.01 | 0.1 | 0.01 m/z or 20 ppm, 0.3 min, 75:25 | No | 2 | | no | yes |
| MSV000082630 | 449 | Q-Exactive | 32000 | 12.5 | | 2E5 | 1E2 | 0.005 m/z or 10 ppm | 0.05 | Baseline cut-off (Min peak height 6E5, Peak dur 0.05-0.4 min, Baseline level 2E5) | 0.01 | 0.1 | 0.005 m/z or 10 ppm, 0.3 min, 75:25 | no | 2 | 0 | no | yes |
| MSV000084007 | 45 | QToF/5600 Triple TOF | | 8 | <2s | 5E2 | 1E1 | 20 ppm | 0.15 | Baseline cut-off (Min peak height 1.8E3, Peak dur 0-3 min, Baseline level 1E3) | 0.02 | 0.15 | 20 ppm, 0.15 min, 75:25 | no | 2 | 2 | no | no |
| MSV000084056 | 9 | Q-ToF/micrO TOF-QII | | 62 | | 3E3 | 5E1 | 20 ppm | 0.1 | Local minimum search (1.0%, 0.1 min, 1.0%, 4000, 1.2, 0.1 - 2.00 min) | 0.02 | 0.1 | 20 ppm, 0.1 min, 75:25 | no | 3 | 2 | | no |
| MSV000084107 | 129 | Q-Exactive plus | 35000 | 18 | | 5E5 | 5E2 | 10 ppm | 0.1 | Local minimum search (30%, 0.1 min, 5.0%, 1500, 2, 0 - 2.00 min) | 0.02 | 0.1 | 7 ppm, 0.2 min, 2:1 | yes | 2 | 2 | no | yes |
| MSV000084063 | 9 | Q-ToF/micrO TOF-QII | 18000 | 25 | | 1E3 | 1E2 | 20 ppm | 0.1 | Local minimum search (1.0%, 0.1 min, 1.0%, 1000, 1.5, 0.1 - 2.00 min) | 0.05 | 0.2 | 20 ppm, 0.5 min, 75:25 | no | 2 | | | no |
| MSV000081832 | 49 | QToF/ Maxis Impact | 17000 | | | 1E3 | 1E2 | 20 ppm | 0.05 | Baseline cut-off (Min peak height 1.0E3, Peak dur 0.01-3 min, Baseline level 1.0E3) | 0.01 | 0.1 | 20 ppm, 0.1 min, 75:25 | no | 2 | 2 | only with MS2 or annotation | yes |
| MSV000084134 | 15 | LTQ-Orbitrap XL | 30000 | 30 | | 3E4 | 1E4 | 10 ppm | 0.1 | Local minimum search (65%, 0.08 min, 0%, 6E4, 1.5, 0-3 min) | 0.2 | 0.2 | 0.002 m/z or 7 ppm, 0.2 min, 3:1 | no | 2 | 0 | only with MS2 or annotation | yes |

### Supplementary Table 5: continued

| ID | Samples | MS Instrument | MS Resolution | Gradient [min] | Duty Cycle Time | $MS^1$ threshold | $MS^2$ Threshold | MS Tolerance | RT Tolerance [min] | Chromatogram Deconvolution Algorithm | $MS^2$ Paring *m/z* Tolerance | $MS^2$ Paring RT Tolerance [min] | Feature Aligment (m/z Tolerance, RT Tolerance, m/z : RT Weights) | Duplicate Row Filter | Min. Peaks in a Row | Min. Isotope Pattern Peaks | $MS^2$ Filtered | Gap Filled |
|---|---|---|---|---|---|---|---|---|---|---|---|---|---|---|---|---|---|---|
| MSV000084148 | 14 | LTQ-Orbitrap XL | 30000 | 30 | | 1E4 | 2E4 | 10 ppm | 0.1 | Local minimum search (85%, 0.1 min, 0%, 2E4, 1.4, 0-2 min) | 0.2 | 0.08 | 0.0015 m/z or 8 ppm, 0.1 min, 3:1 | yes | 3 | 0 | only with MS2 or annotation | yes |
| MSV000084158 | 13 | LTQ-Orbitrap Elite | 120000 | 20 | <2s | 1E4 | 1E2 | 10 ppm | 0.15 | Local minimum search (5%, 0.2 min, 5%, 2E4, 1, 0.02-5 min) | 0.01 | 0.15 | 10 ppm, 0.2 min, 75:25 | no | 2 | 0 | MS2 filter | yes |
| MSV000084170 | 88 | QToF/ Maxis Impact | 17000 | 8 | | 8E2 | 8E1 | 20 ppm | 0.1 | Local minimum search (98.5%, 0.08 min, 0%, 6E3, 1, 0.03-1.5 min) | 0.03 | 0.1 | 0.01 m/z or 20 ppm, 0.1 min, 75:25 | yes | 1 | 2 | no | yes |
| MSV000084118 | 9 | Q-Exactive | 17000 | 10 | <1sec | 1E5 | 1E3 | 10 ppm | 0.15 | Local minimum search (5%, 0.2 min, 5%, 2E4, 1, 0.02-5 min) | 0.01 | 0.15 | 10 ppm, 0.2 min, 75:25 | no | 2 | 0 | MS2 filter | yes |
